# Independent evolution of influenza A virus H1N1 in pigs in Guatemala

**DOI:** 10.1101/2022.07.21.501070

**Authors:** Lucia Ortiz, Ginger Geiger, Lucas Ferreri, David Moran, Danilo Alvarez, Ana Silvia Gonzalez-Reiche, Dione Mendez, Daniela Rajao, Celia Cordon-Rosales, Daniel R. Perez

## Abstract

Commercial swine farms provide unique systems for interspecies transmission of influenza A viruses (FLUAVs) at the animal-human interface. Bidirectional transmission of FLUAVs between pigs and humans plays a significant role in the generation of novel strains that become established in the new host population. Active FLUAV surveillance was conducted for two years on a commercial pig farm in Southern Guatemala with no history of FLUAV vaccination. Nasal swabs (n=2,094) from fattening pigs (6 to 24 weeks old) with respiratory signs were collected from May 2016 to February 2018. Swabs were screened for FLUAV by RRT-PCR and full virus genomes of FLUAV-positive swabs were sequenced by next-generation sequencing (NGS). FLUAV prevalence was 12.0% (95% CI: 10.6% – 13.4%) with two distinct periods of high infection. All samples were identified as FLUAVs of the H1N1 subtype within the H1 swine clade 1A.3.3.2 and whose ancestors are the human origin 2009 H1N1 influenza pandemic virus (H1N1 pdm09). Compared to the prototypic reference segment sequence, 10 amino acid signatures were observed on relevant antigenic sites on the hemagglutinin. We also found that Guatemalan swine-origin FLUAVs show independent evolution from other H1N1 pdm09 FLUAVs circulating in Central America. The zoonotic risk of these viruses remains unknown, but strongly calls for continued FLUAV surveillance in pigs in Guatemala.

**Importance:** Despite increased surveillance efforts, the epidemiology of FLUAVs circulating in swine in Latin America remains understudied. For instance, the 2009 H1N1 influenza pandemic strain (H1N1 pdm09) emerged in Mexico, but its circulation remained undetected in pigs. In Central America, Guatemala is the country with the largest swine industry. We found a unique group of H1N1 pdm09 sequences that suggests independent evolution from similar viruses circulating in Central America. These viruses may represent the establishment of a novel genetic lineage with the potential to reassort with other co-circulating viruses, and whose zoonotic risk remains to be determined.

## INTRODUCTION

Influenza A viruses (FLUAV) infect a wide range of avian and mammalian hosts, including humans. The virus genome is composed of 8 segments of negative single-stranded RNA corresponding to 6 internal (PB2, PB1, PA, NP, M, and NS) and 2 surface (HA and NA) gene segments. Zoonotic FLUAV infections are relatively common and it is accepted that influenza pandemics have resulted from zoonotic FLUAV strains (1). FLUAV infections in swine have a significant economic impact on swine production due to losses caused by the disease. Swine farms present high animal density and provide an environment for close contact between animals and humans. Such environments facilitate the initial spillover between species and transmission (2). Interspecies transmission events of FLUAVs between humans and pigs play a significant role in the generation of novel reassortant strains that transmit among humans and/or swine populations (3). Recent studies suggest that the introduction of human-origin FLUAVs into pigs is a major driver in the independent evolution of FLUAV lineages depending on geographic location (4). The emergence of the 2009 H1N1 influenza pandemic virus (H1N1 pdm09) in Latin America with potential undetected circulation in pigs for several years undetected (5), highlights the need to improve influenza surveillance in these understudied regions to timely detect strains of zoonotic and/or pandemic concern.

In Central America, Guatemala is the country with the largest swine industry. Most of the swine production in Guatemala occurs in farrow-to-finish systems. According to the last census in 2021 by the Ministry of Agriculture, Livestock, and Food of Guatemala, the swine population was estimated to be 294,479 pigs, of which 32,518 were breeding stock (6). Circulation of FLUAV of human origin has been documented previously in swine populations in Guatemala (7); however, only limited genetic data is available. In this study we conducted active surveillance to better understand the epidemiology of FLUAV in swine populations in commercial pig farms in Guatemala (in the absence of vaccination). Our results show that since its introduction in 2009, pdm09-like viruses disseminated and became the predominant subtype in Guatemalan pigs at the time of sampling (2016-2018). Phylogenetic analyses revealed that these viruses cluster with FLUAVs that circulated in 2009 and share nucleotide identities of >97%, representing a unique group of FLUAVs whose zoonotic risk remains to be determined.

## Materials and methods

### Collection site

We selected one commercial farrow-to-finish farm located in Palin, Escuintla (Fig 1A) with 13,044 pigs, and of these 1,233 were breeding stock at the time of the study. The farm production system is divided in boars, sows, replacements, piglets, and growing areas, located on different premises within the farm. Artificial insemination, gestation, and farrowing are performed routinely in the farm. The site did not routinely vaccinate against influenza at the time of the study. Boars, sows, and replacements are in the same facility shared with quarantine, insemination, gestation, replacements, and maternity areas. Piglets are weaned at four weeks of age. After weaning, pigs are moved to a different area where they spend 6 weeks (from week four to week ten). Afterwards pigs are moved to finish stage, where they spend approximately twelve more weeks (from week ten to week twenty-two) to get ready for market. The production cycle of each pig takes approximately 22 weeks. For the past 10 years, the farm reported history of FLUAV exposure in swine.

**Fig 1.**
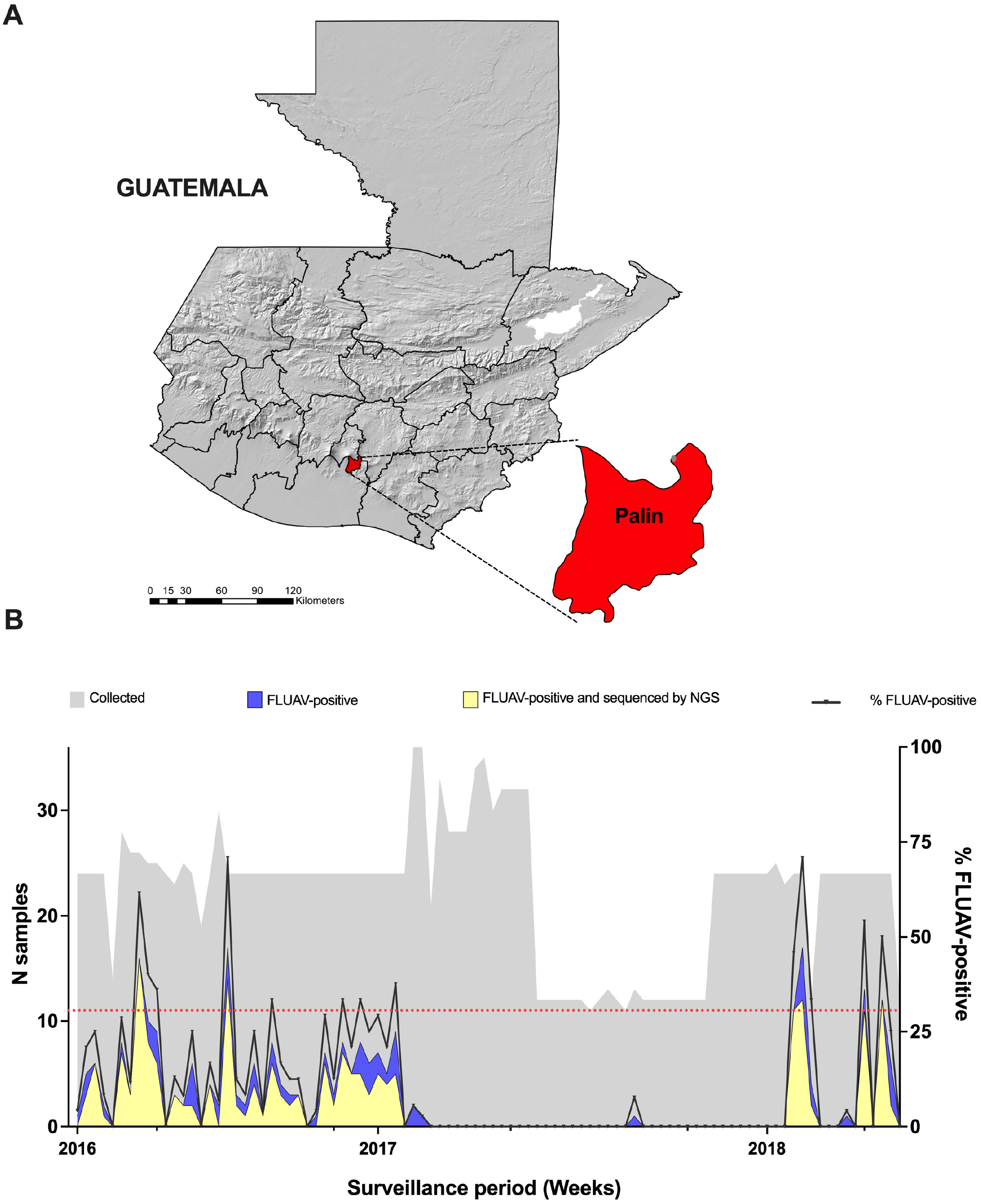
**A) Location of study site in Palin, Guatemala for swine FLUAV surveillance**. Sampled site represents the largest swine-producing region in the country with history of FLUAV circulation. **B) Number of collected samples, and FLUAV-positive, and sequenced samples per week during a two-year period (2016-2018)**. Red dotted line indicates the minimum number of samples required per week to achieve the desired sample size. Two high infection periods were detected by RRT-PCR one from May 2016 to February 2017, and another one from January 2018 to April 2018.

### Sample collection

A total of 2,094 nasal swabs from fattening pigs (6 to 24 weeks old) and sows with respiratory signs were collected from May 2016 to April 2018. Pigs were identified and sampled by farm workers when presenting clinical signs of respiratory disease, including coughing, sneezing, nasal and/or ocular discharges. The farm was sampled twice per week. A minimum of 32 nasal swabs were collected monthly to detect at least 1 positive sample based on a prevalence of 10% or higher, with a 95% confidence and in a group size ≥ 600 (8). Pigs were tattooed with a unique identifier to avoid resampling. Rectal temperature, sex, respiratory signs, and pen density of each animal was recorded at the time of sampling. Nasal swabs were collected and preserved in mL of virus transport medium (VTM) with antibiotics and antimycotics as described previously (7). Sampling of animals was conducted under approved animal use protocols from the MAGA, and the protocols were reviewed and approved by the Institutional Animal Use and Care Committee of University del Valle de Guatemala.

### RNA extraction

Virus RNA from nasal swabs was extracted from 50 µL of supernatant using the MagMAX-96 AI/ND Virus RNA Isolation Kit (Ambion, Austin, TX) according to the manufacturer’s instructions. All extracted RNA was stored at −70°C until use.

### FLUAV detection by RRT-PCR

Swabs were screened in duplicate for FLUAV by real-time reverse-transcriptase polymerase chain reaction (RRT-PCR) targeting the matrix gene, as described previously (7, 9, 10). Briefly, QuantiTect Probe RT-PCR (reverse transcription polymerase chain reaction) Kit (QIAGEN, Hilden, Germany) was used to perform RRT-PCR reactions in the ABI 7300 Real-Time PCR System (Applied Biosystems, Foster City, CA). Each reaction contained 12.5 µL of kit-supplied 2X RT-PCR Master mix, 10 pmol of each primer, 0.3 μM probe, 0.25 µL of kit-supplied enzyme mix, 6.5U RNase inhibitor and 8 µL of RNA template. Thermal cycling conditions were as follows: one cycle of reverse transcription at 50ºC for 30 min and 94ºC for 15 min, followed by 45 cycles of denaturation at 94ºC for 1s and combined annealing and extension at 60ºC for 27s.

### Multi-Segment amplification (MS-RTPCR)

Multi-Segment amplification of FLUAV genes (MS-RTPCR) was performed from RNA extracted from all FLUAV-positive swabs, as described previously (11) with minor modifications. Briefly, 2.5 uL of extracted RNA were used as a template in a 25 uL MS-RTPCR reaction (Superscript III high-fidelity RT-PCR kit, ThermoFisher); using Opti1-F1 (0.06 uM), Opti1-F2 (0.14 uM) and Opti1-R1 (0.2 uM) primers (12). The cycling conditions were as: 55°C for 2 minutes, 42°C for 1 hour, 5 cycles (94°C for 30 seconds, 44°C for 30 seconds, 68°C for 3 minutes), followed by 31 cycles (94°C for 30 seconds, 57°C for 30 seconds, 68°C for 3 minutes). Final extension at 68°C for 10 minutes. MS-RTPCR final product was analyzed in 1% agarose gel to corroborate whole genome amplification.

### Sequencing and genome assembly

MS-RTPCR products were sequenced using the Illumina platform as described previously (12) with minor modifications. Briefly, amplicons from MS-RTPCR reactions were cleaned by 0.45X of Agencourt AMPure XP Magnetic Beads (Beckman Coulter) according to the manufacturer’s protocol and eluted in 30 uL of HyClone molecular biology water (Genesee Scientific). Amplicons were quantified using Qubit buffer kit (Fisher Scientific) in Qubit 3.0 fluorometer (ThermoFisher) and normalized to 0.2 ng/ul. Adaptors were added by tagmentation using the Nextera XT DNA library preparation kit (Illumina). The reaction was set as 60% of the suggested final volume. Samples were purified using 0.7X of Agencourt AMPure XP Magnetic Beads and analyzed on a Bioanalyzer using High Sensitivity DNA kit (Agilent) to determine the distribution of fragment size. Libraries were pooled and normalized to 1-4nM. After denaturation, the final loading concentration of the pooled libraries was 14pM. Libraries were sequenced using the MiSeq Reagent Kit V2 300 cycles (Illumina). Genome assembly was performed using a customized pipeline developed at the Icahn School of Medicine at Mount Sinai Genome assembly was performed using a pipeline previously described (5).

### Minor variants detection

Variant calling was performed as described previously (13) using LoFreq 2.1.3.1

(14) following the Genome Analysis Toolkit best practices (15).

### Phylogenetic analysis

Independent phylogenetic analyses were performed for the surface genes (HA and NA) and internal genes (PB2, PB1, PA, NS, NP, and MP) of the sequenced viruses. The Additional FLUAV genome sequences from human- and swine-origin H1N1 pdm09-like viruses (from 2009 to 2019) from the Americas and avian-origin were downloaded from the Global Initiative on Sharing All Influenza Data (GISAID, http://platform.gisaid.org) and the Influenza Research Database (IRD, http://www.fludb.org). Sequences were aligned with MUSCLE 5.1 (16) and manually trimmed to keep the open reading frame (ORF) of the gene of interest. Background sequences were subsampled to remove identical sequences and those sequences with less than 80% of the total length of the ORF using SeqKit 0.16.1 bioinformatic tool. Representative sequences were randomly selected by region and host. Identical sequences obtained from this study were excluded and only one representative of each sequence was used. Best hit blasts were included in the final tree. The best-fit model of nucleotide substitution was determined for each gene using the Bayesian information criterion (BIC) obtained using jModelTest 2.1.10 (17). For the internal genes, phylogenetic trees were constructed using maximum-likelihood (ML) inference methods, using RAxML 8.2.12 (18) under the general time reversible GTR+G+I nucleotide substitution model with 1,000 bootstrap replicates. Trees were run at least twice to confirm topology. Surface gene (H1 and N1) phylogenies were inferred by time-scaled phylogenetic analysis using the Bayesian Markov chain Monte Carlo (MCMC) approach as implemented in BEAST 1.10.4. A relaxed uncorrelated lognormal (UCLN) molecular clock was used, with a constant population size, and a general-time reversible (GTR) model of nucleotide substitution with gamma-distributed rate variation among sites. Three independent analyses of 50-100 million generations were performed to ensure convergence, sampling every 5,000 states. The burn-in percentage of each dataset was identified using Tracer v1.7.1 and after its removal, results were combined using LogCombiner v1.10.4. The MCC trees were annotated using TreeAnnotator v1.10.4. Outlier sequences were identified using TemEst V1.5.3. When necessary, outlier sequences were removed, and the datasets were run again as described above. Nucleotide alignment, background sequences subsampling, BIC, and phylogenetic analyses (RaxML and Beast) were performed using the computational resources Sapelo2 available at The University of Georgia. The resulting trees were visualized in Figtree 1.4.4 and aesthetically modified using Inkscape v0.48.1 (https://inkscape.org).

### Amino acid sequence analysis

Nucleotide sequences were translated using SeqKit 0.16.1 bioinformatic tool and visualized on Geneious Prime 2021.2.2. The resulting amino acid sequences were compared to prototypic reference sequences to calculate the frequency of phenotypic markers for mammalian transmission/pathogenicity and identify specific motifs in the sequence.

### Statistical analysis

The percentages of positive pigs detected by RRT-PCR were calculated by week, as the total number of positive detected by RRT-PCR by the total number of collected samples per week. To analyze the risk of IAV infection, odds ratios (ORs) were estimated using Stata 15.1 GraphPad Prism v.9.4.0

## RESULTS

### Association between FLUAV infection and male weaning pigs with fever

From May 2016 to April 2018, we performed active FLUAV surveillance in a commercial farrow-to-finish farm located in Palin, Escuintla, ∼60 miles away from the Southern Pacific coast of Guatemala (Fig 1A).

The survey included 31,093 pigs associated to 3,113 cases of respiratory disease. Nasal swabs tested positive for FLUAV by RRT-PCR in 251 out 2,094 samples; with an estimated prevalence of 12.0% (95% CI: 10.6% – 13.4%). Two periods of high FLUAV infection were detected, one from May 2016 to February 2017 and another from January 2018 to April 2018 with prevalence of 18.7% (95% CI: 16.4% – 21.2%) and 17.8% (95% CI: 14.2% – 22.1%), respectively (Fig 1B). Between these two periods, only one FLUAV-positive sample was detected in September 2017. The mean age of the pigs in the farm was 10.5 ± 6.6 weeks. FLUAV was detected in pigs as young as 5 weeks, and as old as 21 weeks. The mean age of FLUAV-positive pigs was 8.0 ± 2.4 weeks. Factors such as age [weaning pigs OR95%=4.1 (1.3, 20.7)], sex [male OR95%=1.4 (1.0, 1.8)] and rectal temperature [fever OR95%=2.7 (2.0, 3.6)] were associated with FLUAV positivity (Table 1). Borderline association was found for animal density, 201-300 animals per pen [OR95%=2.0 (1.0, 5.0)] and FLUAV detection. All animals presented coughing at the time of sampling.

**Table 1.** Risk factors associated with FLUAV detection by RRT-PCR in sampled pigs in Guatemala (2016-2018).

| Characteristic |  | FLUAV positive<br>(%)<br>(n=251) |  | FLUAV negative<br>(%)<br>n=(1,843) |  | OR (95% IC) | P-value |
| --- | --- | --- | --- | --- | --- | --- | --- |
| Sex | Female | 103 | (41.0) | 894 | (48.5) | Referent |  |
|  | Male | 148 | (59.0) | 949 | (51.5) | **1.4 (1.0,1.8)** | **0.0262** |
| Age | Weaning (4-10wks) | 232 | (92.4) | 1236 | (67.1) | **4.1 (1.3,20.7)** | **0.0098** |
|  | Juvenile (11-17wks) | 11 | (4.4) | 296 | (16.1) | 0.8 (0.2,4.7) | 0.7618 |
|  | Semi-adult (18-20wks) | 5 | (2.0) | 245 | (13.3) | 0.4 (0.1,3.0) | 0.2695 |
|  | Adult (>20 wks) | 3 | (1.2) | 66 | (3.5) | Referent |  |
| Rectal temperature (a) | <39.7b | 162 | (64.5) | 1527 | (82.9) | Referent |  |
|  | >39.8 | 89 | (35.5) | 316 | (17.1) | **2.7 (2.0,3.6)** | **0.0000** |
| Density<br>(animals/pen) | <100 | 8 | (3.2) | 100 | (5.4) | Referent |  |
|  | 101-200 | 24 | (9.6) | 315 | (17.1) | 1.0 (0.4,2.5) | 0.9084 |
|  | 201-300 | 110 | (43.8) | 672 | (36.5) | 2.0 (1.0,5.0) | 0.0558 |
|  | 301-400 | 107 | (42.6) | 703 | (38.1) | 1.9 (0.9,4.7) | 0.0871 |
|  | >400 | 2 | (0.8) | 53 | (2.9) | 0.5 (0.0,2.5) | 0.3428 |

<sup>a</sup>: Normal temperature ranges between 38.7–39.7. OR: odds ratio.

### Swine-origin FLUAVs in Guatemala show independent evolution from other H1N1 pdm09 FLUAVs circulating in Central America

From the 251 FLUAV positive samples, 157 (62.5%) amplified at least one gene segment by multi-segment RT-PCR (MS-RT-PCR). These samples were subsequently sequenced by NGS, producing 57 complete genomes (53 from the 2016-17 period and from the 2018 period). The total number of full-length sequences by segment and the number of unique Open Reading Frames (ORF) sequences are shown in (Table 2). To build the phylogenetic trees, only one representative from identical sequences was used resulting in 273 unique nucleotide sequences (225 from the 2016-17 period and 48 from the 2018 period, accession numbers can be found at the Supplemental File 1).

**Table 2.**
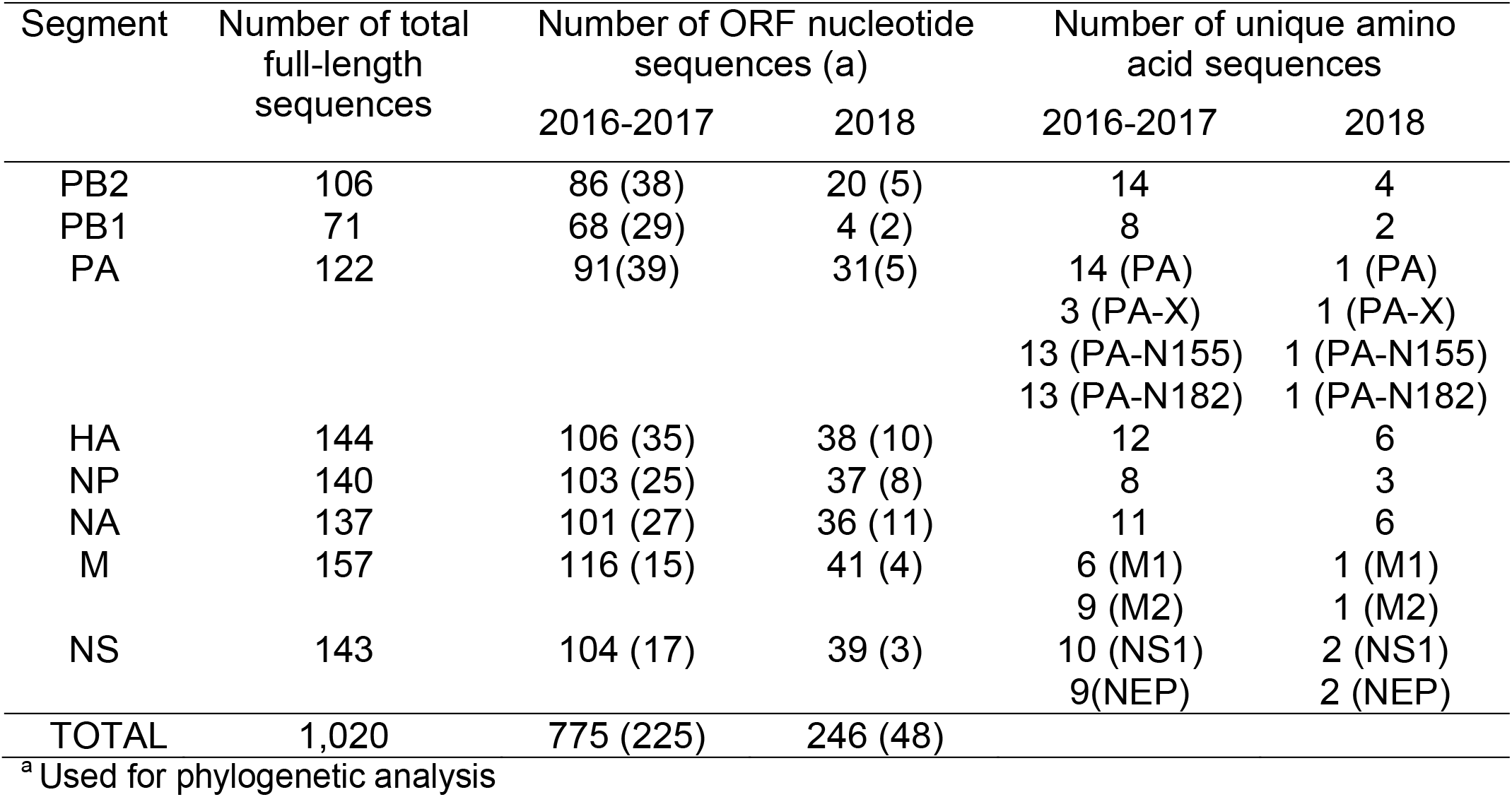
Total number of sequences obtained from sampled pigs in Guatemala (2016-2018).

Phylogenetic analyses of all gene segments show 2 clusters, one from samples collected from May 2016 to February 2017 and the other from samples collected in 2018. Within each cluster, all gene segments showed >99% sequence identity. Both clusters share a common ancestor derived from the H1N1 pdm09 FLUAV lineage. The HA segments of all samples belonged to the FLUAV H1 swine clade 1A.3.3.2 derived from the H1N1 pdm09 FLUAV lineage using the “swine H1 clade classification tool” of the Influenza Research Database (IRD) (19).

The HA and NA phylogenies of the Guatemalan FLUAVs show clear separation from other contemporary human H1N1 pdm09 FLUAVs and all swine FLUAVs detected in Central America whose sequences are available in the IRD and GISAID databases (Fig 2). The HA and NA gene segments of Central American human origin FLUAV isolates from 2009 are the most closely related ancestors of the HA and NA segments of the swine FLUAVs from Central America even though the samples in this study were collected during 2016-2018. It is estimated that these viruses were introduced early during the A(H1N1)pdm09 pandemic (∼2010.8 [2009.4-2012.7 95% HPD] for HA and 2010.4 [2009.1-2012.3 95% HPD] for NA).

**Fig 2.**
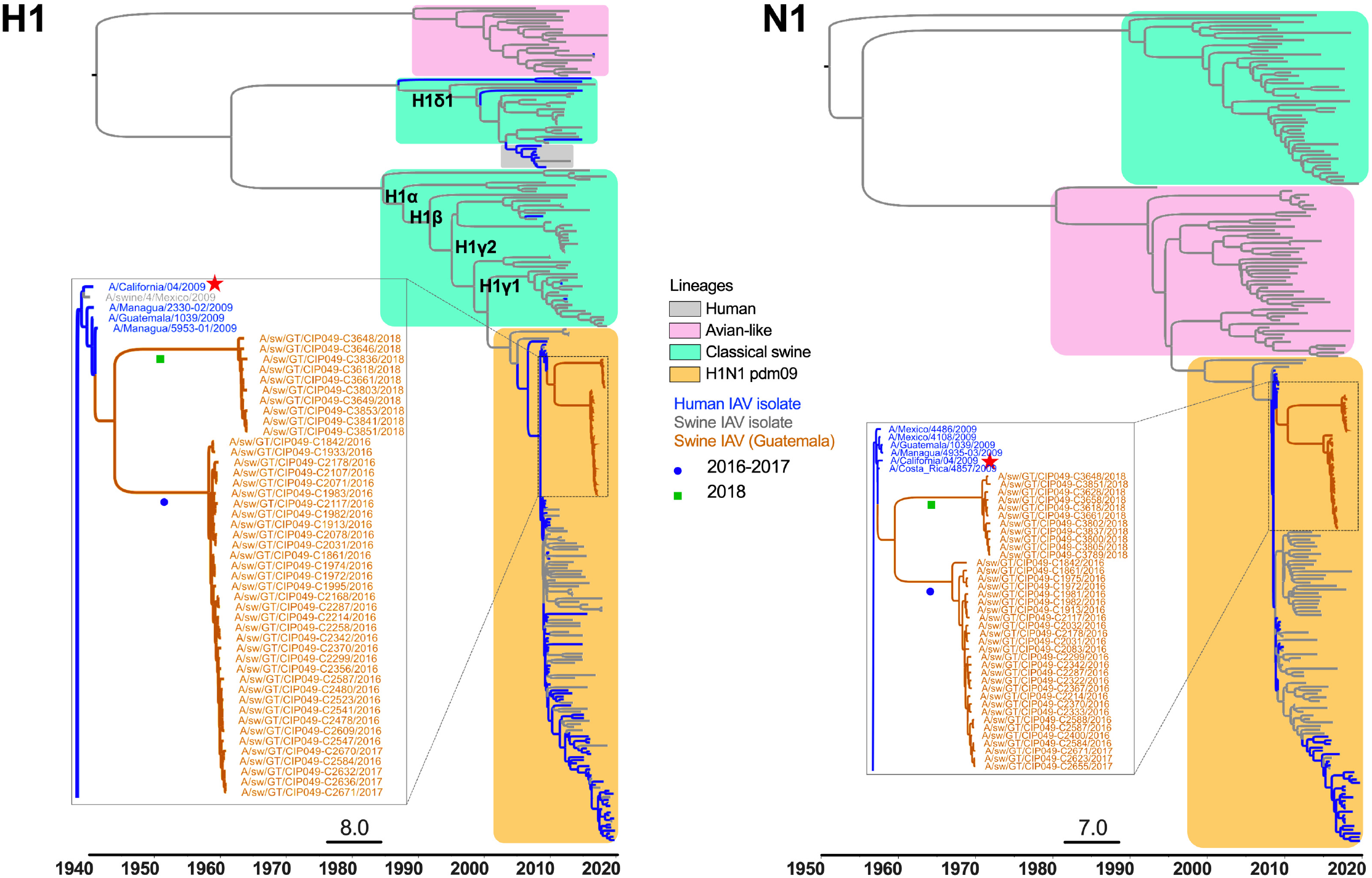
Surface gene segments (H1 and N1) time scaled MCC trees inferred for H1N1 pdm09 from swine and human coding sequences (2008-2019) for 273 (H1) and 321 (N1) sequences. Reference strain Ca04 is highlighted with a star symbol. Identical sequences of Guatemalan viruses were removed. The swine FLUAVs from Guatemala are marked with a blue circle (May 2016 to February 2017 cluster) and a green rectangle (2018 cluster).

Phylogenetic analyses of the polymerase genes (PB2, PB1 and PA) showed similar clustering within H1N1 pdm09 FLUAV identified in humans in 2009 (Fig 3). NP and M1 genes from Guatemalan swine IAVs clustered with FLUAVs circulating in the Americas in swine and humans in 2009 respectively. Interestingly, the NS gene segment was shown in two clusters, those from the 2016-17 period associated with NS sequences of swine FLUAVs identified in Japan in 2017 and those from the 2018 period associated with swine FLUAVs circulating in Thailand between 2013-2017, consistent with possibly two independent introductions (Fig 4).

**Fig 3.**
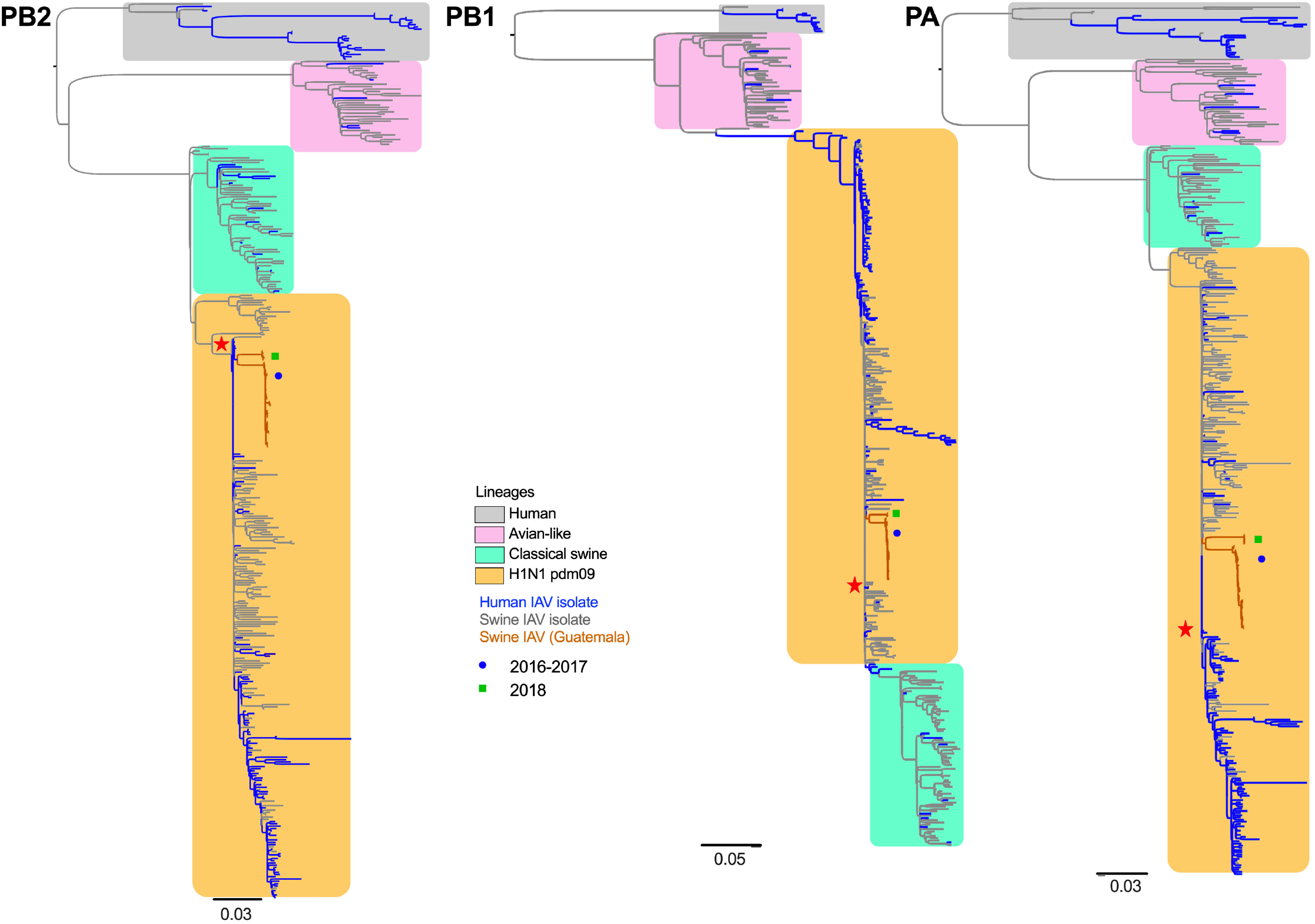
Internal gene segments (PB2, PB1, PA) and phylogenetic inference for H1N1 pdm09 from swine and human coding sequences (2008-2019) for 390 (PB2), 378 (PB1) and 399 (PA) sequences. Maximum Likelihood phylogenetic inference using the best-fit model. Reference strain Ca04 is highlighted with a star symbol. Identical sequences of Guatemalan viruses were removed. The swine viruses from Guatemala are marked with a blue circle (May 2016 to February 2017 cluster) and a green rectangle (2018 cluster).

**Fig 4.**
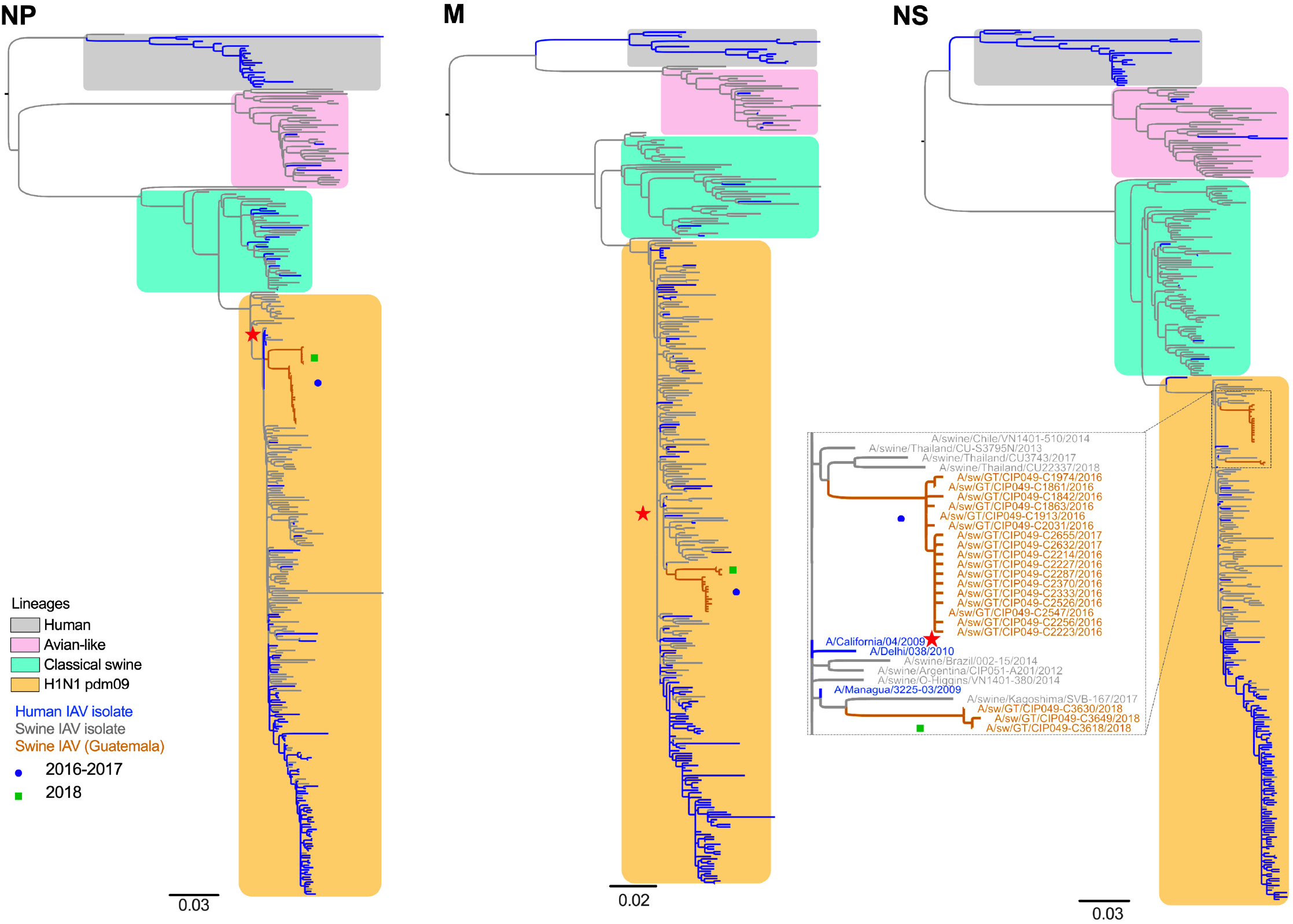
Internal gene segments (NP, M, and NS) and phylogenetic inference for H1N1 pdm09 from swine and human coding sequences (2008-2019) for 360 (NP), 347 (M), and 378 (NS) sequences. Maximum Likelihood phylogenetic inference using the best-fit model. Reference strain Ca04 is highlighted with a star symbol. Identical sequences of Guatemalan viruses were removed. The swine viruses from Guatemala are marked with a blue circle (May 2016 to February 2017 cluster) and a green rectangle (2018 cluster).

### Five amino acid signatures were fixed on relevant antigenic sites on the HA compared to the Ca04 reference sequence

Amino acid differences in HA observed in swine H1N1 pdm09 FLUAVs from Guatemala were compared to the reference strain A/California/04/2009 (H1N1) (Ca04, Fig 5 and Fig S1). The HA ORF encoded the same cleavage site sequence indistinguishable from the Ca04 strain (PSIQSR’GLF). In the rest of the HA ORF, up to 39 amino acid differences were observed compared to the Ca04 reference sequence (Fig S1), 25 of those on the HA1 globular head with 10 falling within antigenic sites Cb (S71F), Ca2 (P137S, H138Y, A141T), Ca1 (G170E, R205K, E235G) and Sb (S185N, S190T, A195S) (H1 numbering, mature protein). Minor variant frequency analysis revealed other differences on the HA1 globular head in samples from different animals (A141E, L161P, T184A, T184I, D187G, Q188R and E235K) with allele frequency ranging from 2.3% to 36.3% in at least 8 viruses from the 2016-2017 period. Most amino acid differences in the HA1 region occurred at high frequency (>0.95) (Fig 6). Five of these amino acid positions were in common among the HA ORFs from the two high FLUAV infection periods (P83S, T197A, R205K in antigenic site Ca1, I321V and D346E). The HA R45Q and the HA R45K were fixed in viruses from the 2016-2017 and 2018 periods, respectively (Fig S1). The HA L32V was the most common signature in viruses from both infection periods (>100 HA sequences), except for one virus each from 2016 and 2018 (L32M and L32I, respectively). S71F (Cb), P137S (Ca2) were fixed in viruses from the 2016-2017 and A141T and S190T in the 2018 period.

**Fig 5.**
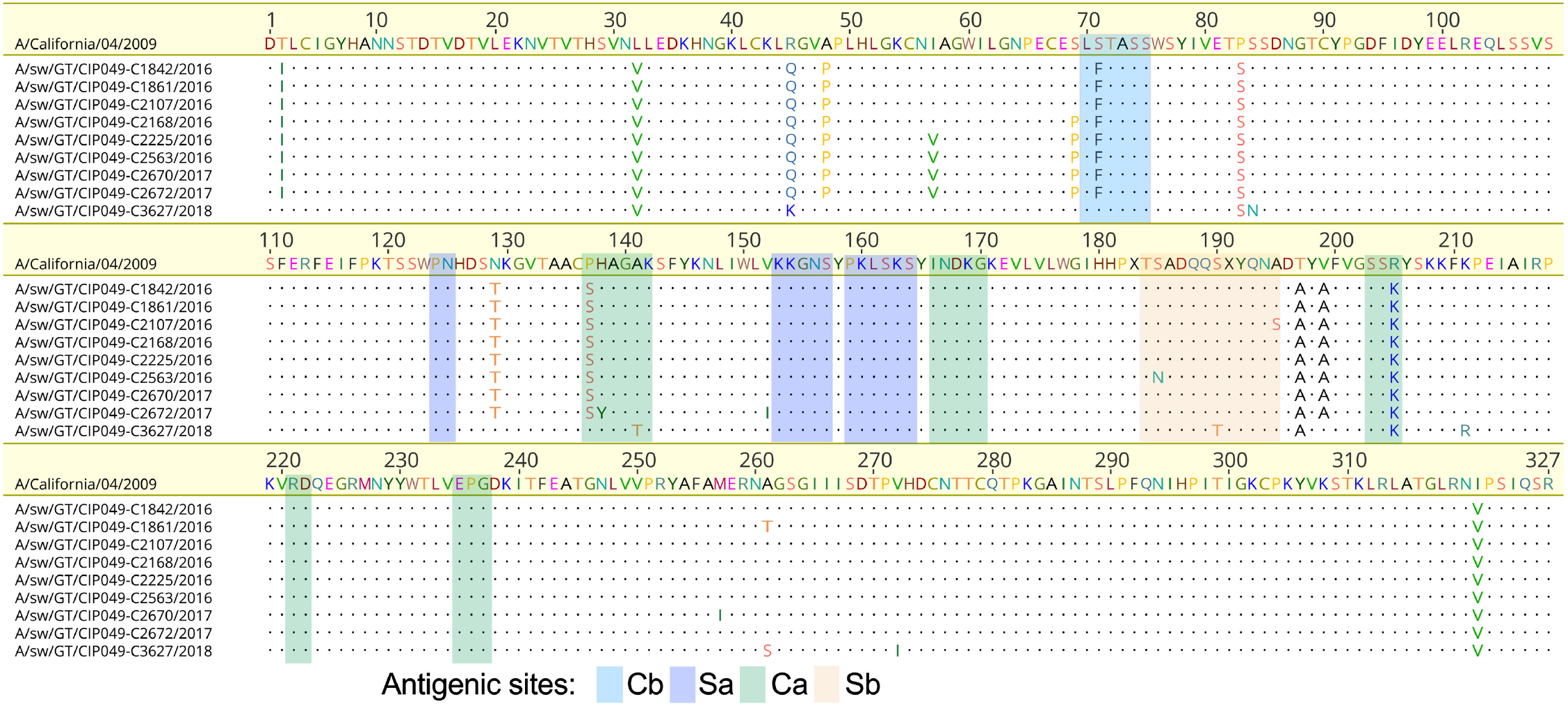
Distribution of amino acid differences in the HA1 domain (H1 numbering without the signal peptide) of unique H1N1 pdm09 swine viruses from Guatemala aligned to Ca04 (GenBank accession no. GQ117044.1). Dots represent amino acids identical to the reference strain. The antigenic sites (Cb, Sa2, Ca1 and Sb) are shown in colored boxes.

**Fig 6.**
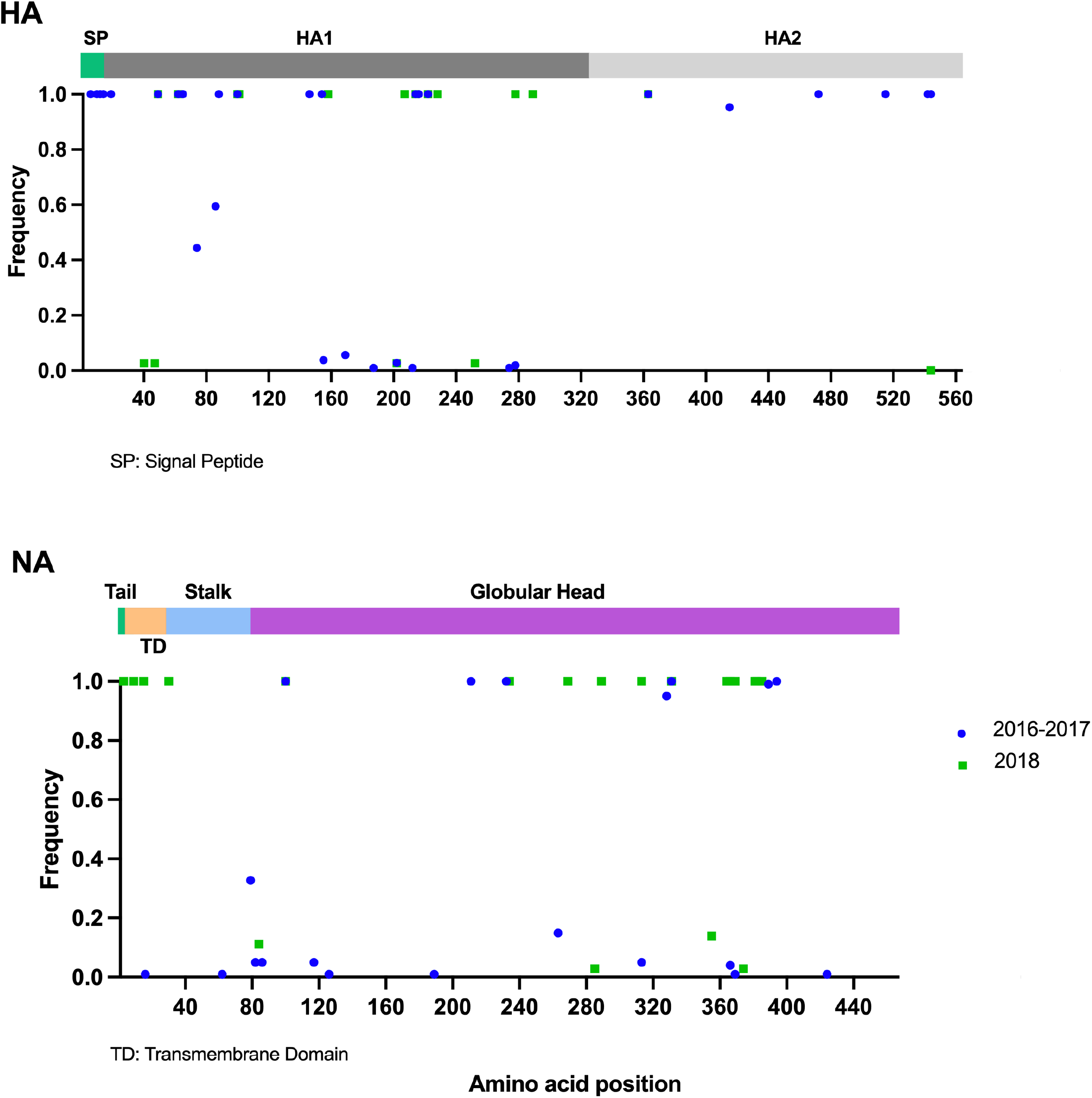
Distribution and frequency of amino acid differences in surface proteins (HA and NA) from H1N1 pdm09 swine viruses from Guatemala in comparison to the reference strain Ca04 (GenBank accession no. GQ117044.1 and MN371610.1). Known protein domains or important sites of pdm09 haplotypes available in public databases are represented by colored blocks. Amino acid position is shown in the x-axis.

### Amino acid signatures elsewhere in predicted ORFs of swine FLUAVs from Guatemala

In the NA ORF, 36 amino acids differences were found compared to the N1 NA from the Ca04 reference strain (Fig S1), but most were found fixed in NA sequences from the 2018 period (18 amino acid signatures in 35 sequences). In contrast, only 5 amino acid signatures were identified in >90% out 101 sequences in the NA sequences from the 2016-2017 period. Only the Y100H signature was in common among sequences between the two periods, whereas the NA K331R and NA K331N were fixed in strains from the 2016-2017 and 2018 periods, respectively. None of these differences were present in either in the catalytic site (118, 151, 152, 224, 276, 292, 371, and 406, N2 numbering) or the framework residues (119, 156, 178, 179, 198, 222, 227, 274, 277, 294, and 425, N2 numbering) (20), and no amino acid changes in known drug binding sites were found (Fig 6).

In the internal gene segments, eight substitutions that appeared in both periods were found in PB1, PA, NP and M2 (Fig 7 and Fig S2-S3). Compared to the Ca04 reference, the PB1 fixed differences I179M, K353R, and N455D are located on predicted RNA-dependent RNA polymerase motifs. PA presented fixed amino acid differences in the NLS region (P224S) and the PB1 binding domain (Y650F), while NP showed an amino acid difference overlapping the PB2 binding and RNA binding domains (D53E). M2 showed two fixed differences found in both periods, located at amino acid positions 55 and 61 (F55I and R61K). All swine origin FLUAV Guatemalan viruses presented a truncated form of the PB1-F2 protein of 11 aa, due to premature stop codons at positions 12 and 58, in common with the Ca04 reference strain (21). All swine origin FLUAV Guatemalan viruses encode the PA-X functional protein (232 aa), as well as the ORFs for PA-N155 (562 aa) and PA-N182 (535 aa). No alternative amino acid differences were found in PA-X in either period compared to Ca04. In contrast, PA-N155 and PA-N182 ORFs show two amino acid differences, P70S/Y498F and P43S/Y469F, respectively (Fig S4).

**Fig 7.**
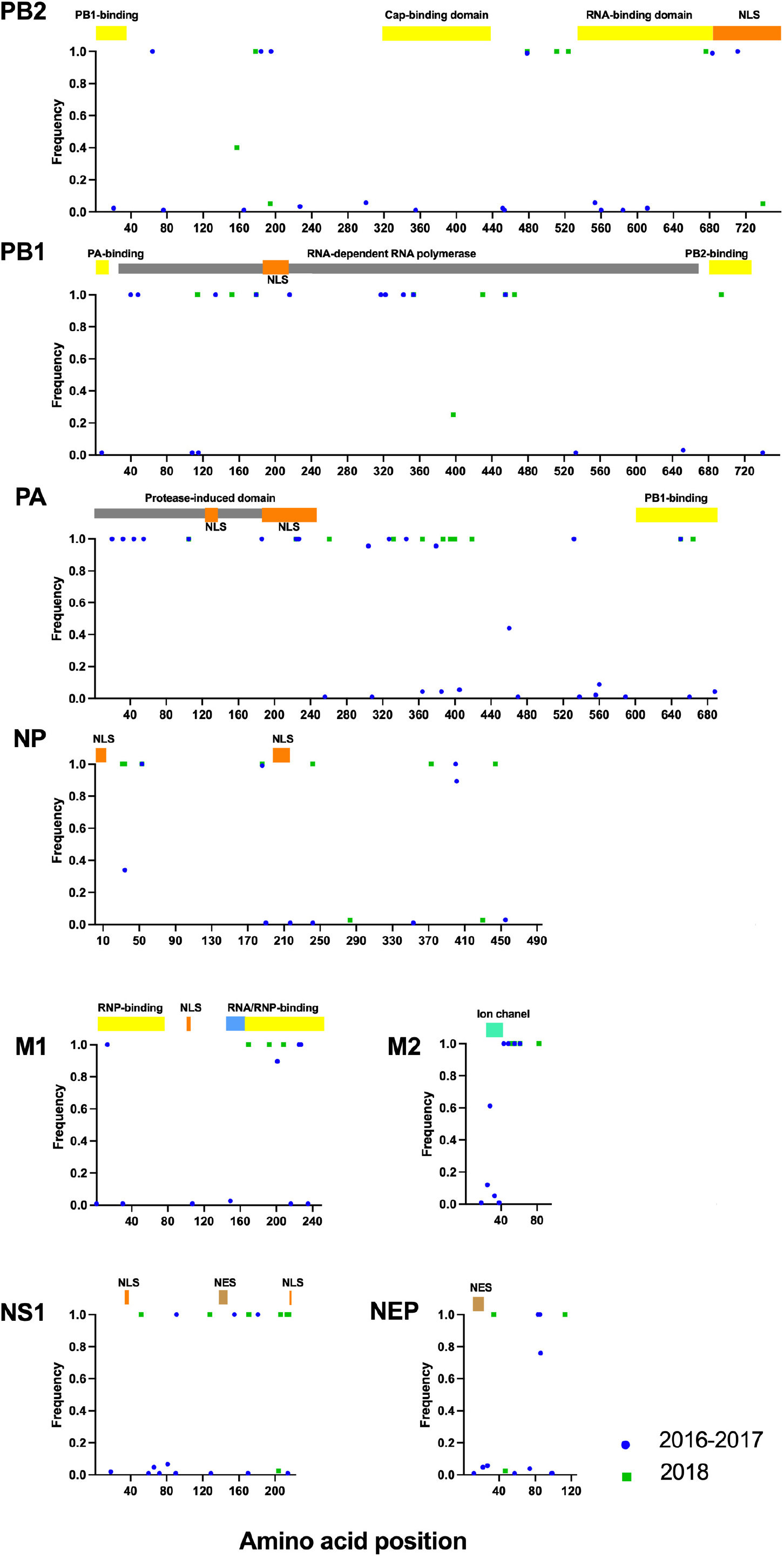
Distribution and frequency of amino acid differences in PB2, PB1, PA, NP, M1, M2, NS1 and NEP from swine H1N1 pdm09 FLUAVs from Guatemala in comparison to the reference strain Ca04 (GenBank accession no. MN371615.1, MN371613.1, MN371611.1, MN371617.1, FJ969513.1, and FJ969514.1). Known protein domains or important sites of pdm09 haplotypes available in public databases are represented by colored blocks.

No mammalian-associated virulence markers in PB2 (E627K, D701N), PB1-F2 (N66S), NS (S42P, D92E and V149A), and M2 (V27A) were found for any of the analyzed sequences. All swine origin FLUAV Guatemalan viruses showed the PA S409N signature predictive of increased virulence and the M2 S31N marker of amantadine resistance like other Eurasian swine lineage M segments (22-24).

## DISCUSSION

Close contact between humans and pigs in swine production systems may result in bidirectional transmission of different FLUAVs between the two species. Systematic surveillance of FLUAV in swine production farms provides a unique opportunity to study how these viruses may jump and adapt at the human-animal interface and identify novel strains that may become established in the new host population. Based on an extensive FLUAV weekly surveillance in pigs from 2016 to 2018, we detected an overall FLUAV prevalence of 12% in animals with respiratory disease, similar to previous studies in pigs in Guatemala (7). It must be noted that sick animals were identified and sampled based on observation of coughing (regardless of other signs), which may have resulted in underrepresentation of sampling and consequent underestimation of the FLUAV prevalence found in this study. Younger pigs showed the highest positivity rate of FLUAV and were associated with higher rates of infection, consistent with previous surveillance studies (7, 25). Factors such as sex (males) and fever were also associated with FLUAV positivity.

During the span of 2 years, we detected two high infection periods but not a seasonality pattern. Only the H1N1 subtype from the H1N1 pdm09 lineage was identified, despite other FLUAV subtypes being reported in humans and swine in Guatemala and Central America at the time of the surveillance period (7, 26). National records from Guatemala indicate minimal to zero circulation of H1N1 pdm09 FLUAVs in humans at the time of pig sampling for this study. This observation suggests a prior introduction of H1N1 pdm09 FLUAVs into pigs that remained endemic in the swine population in Guatemala. Interestingly, a sharp reduction in FLUAV-positive samples was observed for approximately 12 months, during most of 2017. This drop in detection could be explained by an increase in prevention and biosafety practices following the first year of the study (annual seasonal influenza vaccination of workers; quarantine of sick pigs; use of hand disinfectant and mask respirators by workers; and enforcement of the shower-in/shower-out procedures), as reported by farm owners. However, the reason for the subsequent increase in detections seen in 2018 remains unknown, although it is consistent with the notion of introduction of a virus strain through replacement animals brought into the farm.

We obtained sequences from >75% of the FLUAV-positive samples and nearly 25% of complete genomes by NGS directly from the original swabs. This methodology allowed us to improve the number of characterized samples and reduce potential selection bias introduced by virus isolation prior to sequencing (12). Interestingly, all samples presented high number of defective RNAs, as it was observed in agarose gels and later by the valley-like shape in the graphs of the polymerase genes (PB2, PB1, and PA) of the NGS coverage read maps (Fig S5). These particles are truncated forms of FLUAV generated by most viruses during virus replication that retain the terminal sequences necessary for virus packaging (27) and are mostly present in the polymerase segments (28, 29). The function is not fully understood; but it is hypothesized that they could play a role in maintaining low levels of replication of infectious virus (30).

Phylogenetic analyses showed that the swine origin FLUAV Guatemalan viruses are clearly segregated from other H1N1 pdm09 viruses in the Americas, suggesting independent evolution of these viruses after introduction and subsequent circulation in pigs, consistent with reports in other regions (31). At least two independent introductions were observed as noted with the separated clusters of Guatemalan samples. Interestingly, the viruses seemed to have been introduced in 2010 during the H1N1 pdm09 pandemic and persisted as separate clades for ∼6-8 years before detected by our surveillance. Long persistence sustained prevalence of human-origin FLUAVs in environments with significant interaction between two FLUAV hosts might lead to the generation of strains of pandemic concern.

Little human seasonal influenza sequence data is available from Guatemala and Central America and much less swine FLUAV sequence data exists. This gap in part explains the results of the phylogenetic relationships with the most closely related ancestors. Independent virus evolution events have been documented previously in pigs, particularly after introduction of human FLUAVs, showing long phylogenetic branches between the swine strains and their putative human FLUAV ancestor, as shown previously from other Latin American countries (4, 32). Here, we describe the circulation and evolution of H1N1 pdm09 lineage viruses in Guatemala that may represent the establishment of a novel genetic lineage with the potential to reassort with cocirculating viruses. The mutations found in relevant HA1 antigenic sites may lead to differences in antigenic relationships with other H1N1 pdm09 viruses of human- or swine-origin; however, the effect of these mutations and zoonotic risk remains to be determined. These observations highlight the need for increased and sustained influenza surveillance within understudied regions.

## Data availability

Nucleotide genomic sequences from all Guatemalan viruses have been deposited at the NCBI Database under the following accession numbers ON822118-ON823137.

## Acknowledgments

Jorge Paniagua, Silvia Ramírez, field epidemiologists from MAGA (The Ministry of Agriculture, Livestock and Food of Guatemala) and farm owners and workers for allowing sample collection.

## Authors’ contributions

LO: designed studies, processed field samples, performed sequencing, developed methods, analyzed data, and wrote and edited the manuscript. LF: edited the manuscript. GG: performed sequencing and edited the manuscript. AG: edited the manuscript. DM: collected field samples, designed map and edited the manuscript. DA: designed studies and edited the manuscript. CC: designed studies and edited the manuscript. DM: DR: edited the manuscript. DR: edited the manuscript. DP: designed studies, analyzed data, and wrote and edited the manuscript. All authors have read and agreed to the published version of the manuscript.

## Supplemental Material

**Fig S1. HA (H1 numbering) and NA (N2 numbering) unique amino acids of H1N1 pdm09 swine viruses from Guatemala aligned to Ca04 (GenBank accession no. GQ117044.1 and MN371610.1)**. Dots represent amino acids identical to Ca04. The antigenic sites for HA (Cb, Sa2, Ca1 and Sb) are shown with different colors. Viruses identified during May 2016 to February 2017 are shown in blue and those collected in 2018 in green.

**Fig S2. PB2, PB1, PA and NP unique amino acids of H1N1 pdm09 swine viruses from Guatemala aligned to Ca04 (GenBank accession no. MN371615.1, MN371613.1, MN371611.1 and MN371617.1)**. Dots represent amino acids identical to Ca04. Viruses collected during May 2016 to February 2017 are shown in blue and those collected in 2018 in green.

**Fig S3. M1, M2, NS1 and NEP unique amino acids of H1N1 pdm09 swine viruses from Guatemala aligned to Ca04 (GenBank accession no. FJ969513.1, FJ969514.1)**. Dots represent amino acids identical to Ca04. Viruses collected during May 2016 to February 2017 are shown in blue and those collected in 2018 in green.

**Fig S4. PA-X, PA-N155 and PA-N182 unique amino acids of H1N1 pdm09 swine viruses from Guatemala aligned to Ca04 (GenBank accession no. MN371611.1)**. Dots represent amino acids identical to Ca04. Viruses collected during May 2016 to February 2017 are shown in blue and those collected in 2018 in green.

**Fig S5. Coverage plot of sequenced samples**. Gray lines show the coverage distribution from each individual sample. The red line depicts the geometric mean.

